# MembraneBuilder: High-speed membrane construction for large molecular dynamics simulations

**DOI:** 10.1101/2025.11.20.689436

**Authors:** Cayson Hamilton, Gus Hart

**Affiliations:** Brigham Young University

## Abstract

Membrane building and embedding of proteins into a membrane is a crucial step in preparing a simulation for Molecular Dynamics (MD). When an MD simulation contains several million atoms, embedding the protein into a membrane becomes computationally expensive. `MembraneBuilder` [1] speeds up the generation of these membranes by simplifying the lipid insertion process, which allows membranes to be generated for these larger systems. Realistic lipid densities and elimination of infinite forces caused by overlapping atoms are achieved at a low computational cost. See Figure 1 for specific time comparisons. The comparisons emphasize that not only does `MembraneBuilder` run orders of magnitude faster than state-of-the-art programs [2, 3, 4], it enables generation of systems several orders of magnitude larger.

`MembraneBuilder` enables customizable embedding of proteins into a membrane. Users can modify lipid types and their ratios, the location to insert the membrane, the size of the membrane box, as well as the lipid density. The program reduces the computational time required to perform a membrane embedding by bypassing many of the computationally expensive steps, such as identification of polar/non-polar sections of the protein and exhaustive packing of the lipids into the membrane with the intent to avoid steric clashes. Efficient packing begins with constricting the lipid bonds by a given factor in *x, y*, and *z* directions before inserting them on a grid. Grid spacing is determined by the width of the constricted lipids. The grid spacing is smaller than in other lipid-generation tools due to the constriction process. The constricted bonds will relax and the lipids will assume realistic interweaving during energy minimization and equilibration.

## 2 Statement of Need

Molecular dynamics is a method to explore the mechanistic interactions of molecules and proteins and test hypotheses regarding their structure and function. For these systems to accurately represent the environment they simulate, they need to have similar molecular structures. Obviously, this is the case with membrane proteins, which are of significant importance in the field of molecular dynamics. Obtaining accurate lipid ratios, densities, and placement is important to accurately reflect the cellular system.

**Figure 1:**
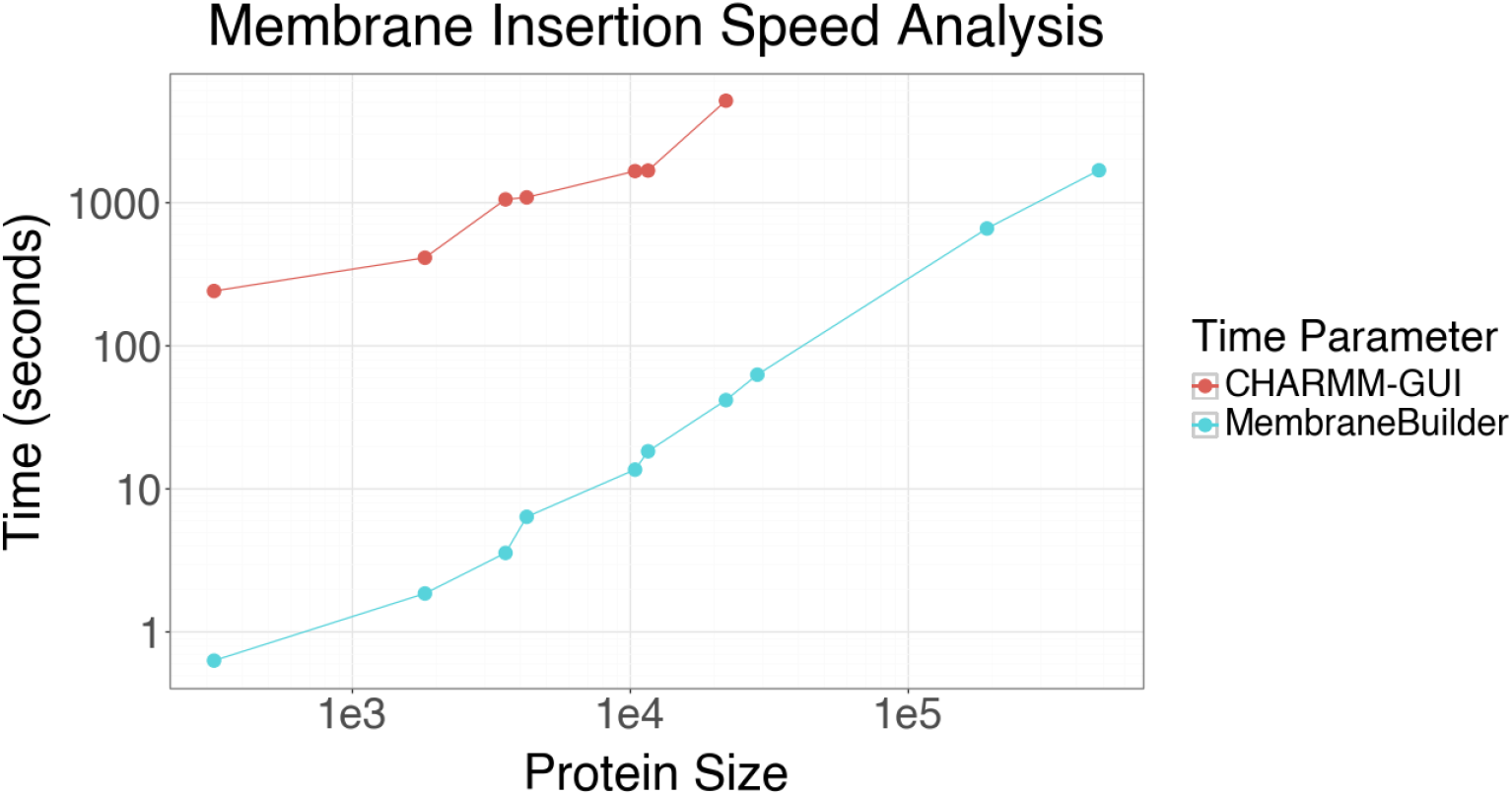
Speed Analysis: Comparison between the amount of time taken to run MembraneBuilder and CHARMM-GUI with varying protein sizes and their requisite amount of lipids. Note that the *x* and *y* scales are on a log scale, which further emphasizes the efficiency of MembraneBuilder.

While standard membrane building tools are effective when working with small systems, they fail when dealing with large systems that require insertion of more lipids. Larger systems cause issues due to packing irregularities or artifacts, incorrect periodic boundary conditions, user interface crashes, or lack of sufficient computational resources. Even if membrane generation is successful, it may take days to weeks to complete and may contain steric clashes making MD impossible. In most cases, large systems cannot even be processed. CHARMM-GUI [2, 3, 4], the most common membrane builder, can only process files smaller than 100 MB or containing less than 3 million atoms [5]. Most importantly, `MembraneBuilder` can process large systems efficiently while avoiding steric clashes. Beyond that, `MembraneBuilder` provides options to customize lipid ratios, densities, and location. We have built a membrane system with 14 million atoms and the simulations of the system run without error. [6].

## 3 Methodology

The core algorithm centers around the idea of compressing atom positions within the lipid to reduce the amount of space each lipid occupies, allowing insertion of more lipids in the same space with no overlapping atoms (avoiding steric clashes). The lipids are constricted by two user-defined factors, one for compression in the *x*-*y* plane and another in *z* direction. An example constriction is shown for the POPE lipid in Figure 2. Panel B shows that the lipid has the same topology, just constrained to a smaller volume, allowing for tighter packing.

**Figure 2:**
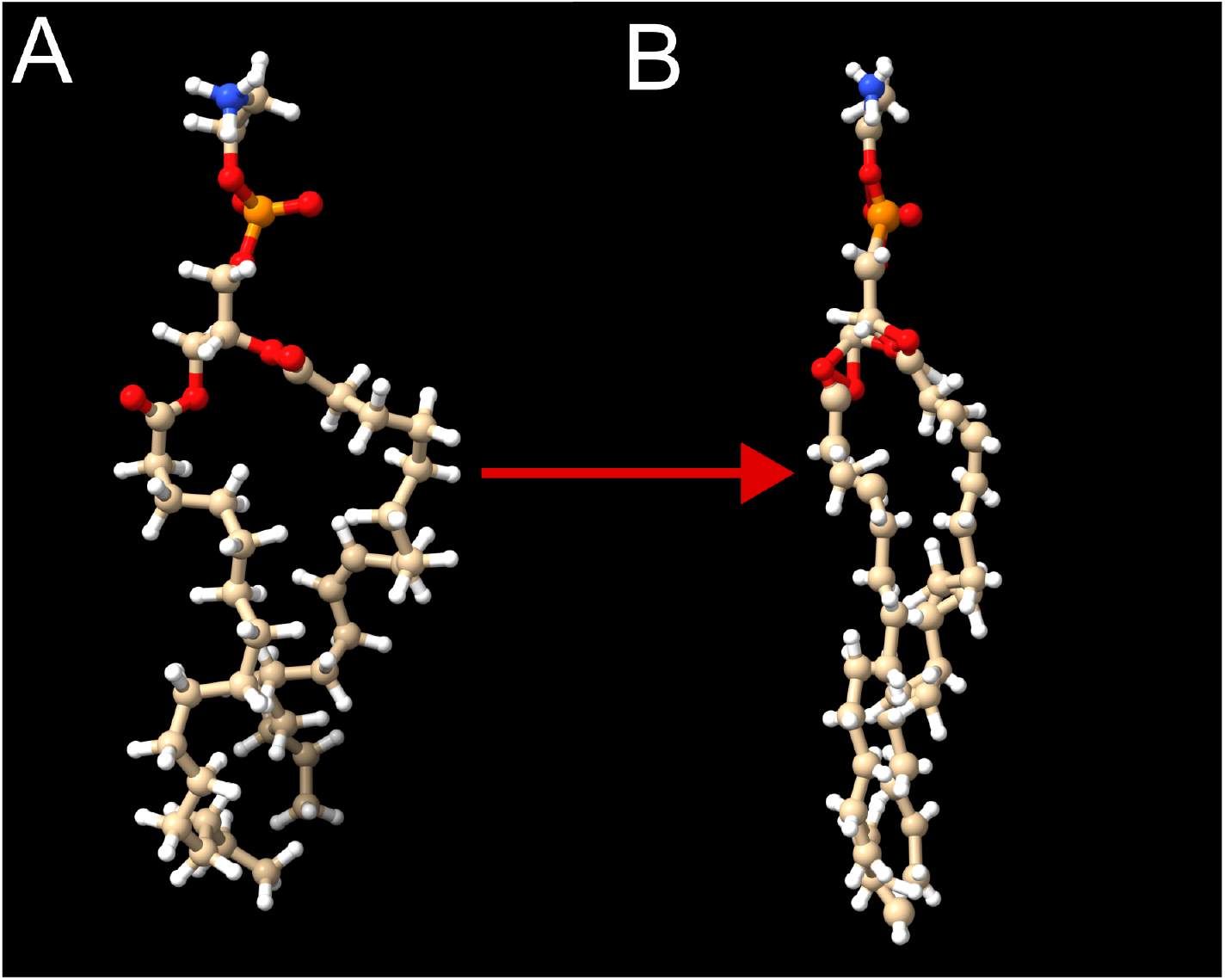
Lipid Constriction: (A) POPE in the unconstricted configuration. (B) POPE constricted by 0.55 in the *x*-*y* directions and stretched by 1.1 in the *z* direction.

In addition to lipid-lipid steric clashes, the algorithm avoids lipid-protein clashes with the membrane protein. The first step is generating a representation of the 3D space the protein occupies. This is done by identifying the maximum and minimum *x* and *y* values at 2 Å intervals moving up the *z*-axis. These computations are done before lipid insertion to reduce computational demand later in the process.

Lipid insertion then begins at the specified *z*-coordinate and at (− [*BoxSize/*2], − [*BoxSize/*2]) in the (*x, y*) plane (the protein must be centered prior to running `MembraneBuilder`). After a lipid is inserted, the insertion moves the exact width of the inserted lipids plus a customizable buffer, which can be adjusted to customize lipid density. Once the insertion reaches the *x* coordinate corresponding to the defined box size, *x* is reset and *y* is adjusted by the maximum width of all the lipids being inserted. This process continues until both *x* and *y* reach the box size. Then, the lipids are reflected across the *x*-*y* plane, moved upwards in the *z*-direction by a specified buffer, and the process repeats to generate the second leaflet. The buffer between lipids can be adjusted to allow for more or less extreme constriction of the lipids while continuing to avoid clashes.

Before any lipid is inserted into the system, the maximum and minimum *x* and *y* protein coordinates are retrieved from the pre-computed values for the positions of the center, highest, and lowest atoms in the lipid. If the coordinates of any of these three spots on the lipid have an *x* or *y* coordinate that is smaller than the maximum value and greater than the minimum value (they overlap the protein), the lipid is not inserted, and insertion moves to the next position. This process greatly reduces likelihood of steric clashes in the generated membrane configuration, and performs the insertion in a very timely manner.

An example insertion can be seen in Figure 3, which was used in research performed on the *Legionella pneumophila* Type IV Secretion System (T4SS) [6]. The lattice pattern is clearly visible, an effect which disappears once energy minimization and equilibration have been run and the system moves towards its natural configuration.

**Figure 3:**
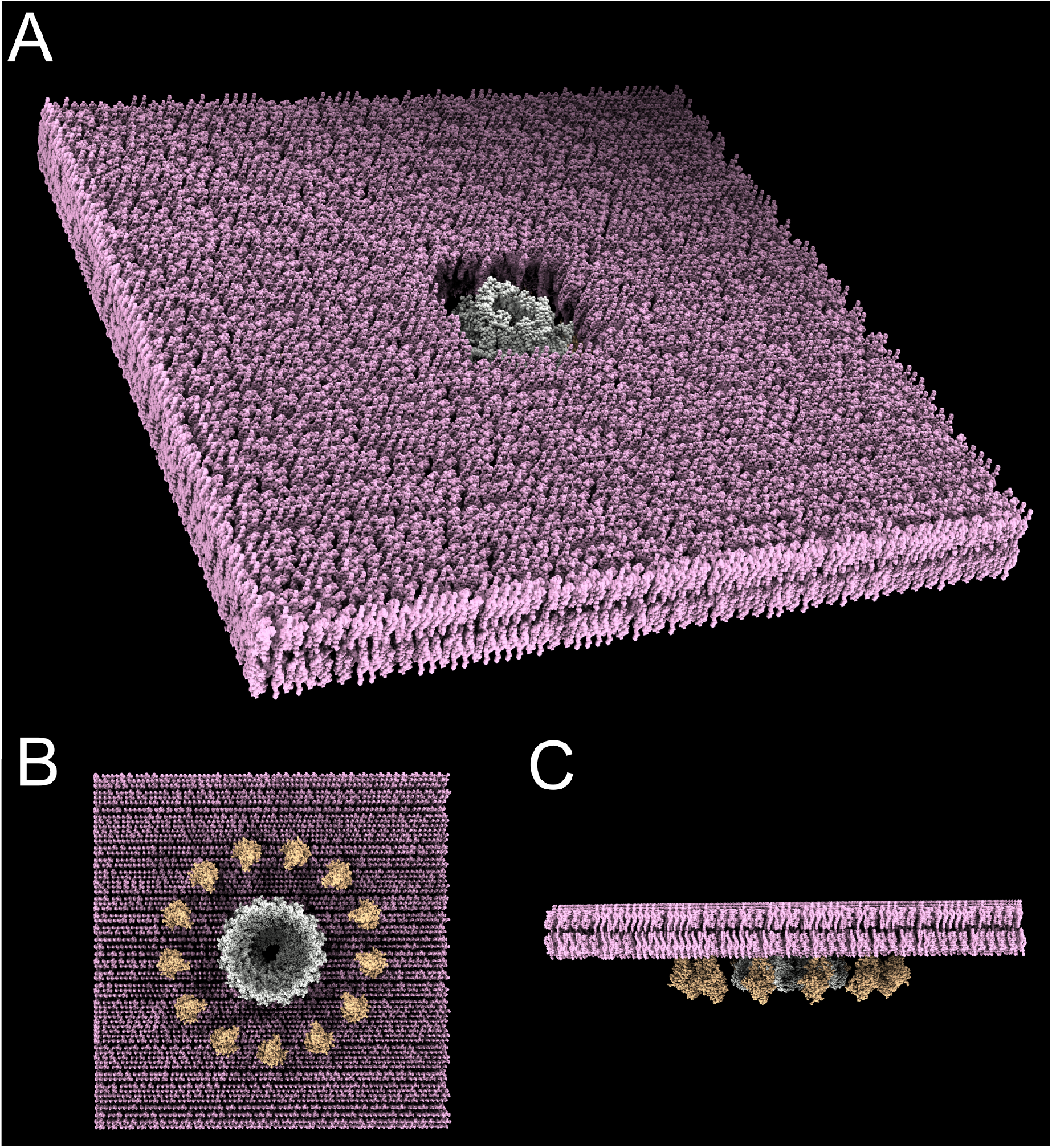
Membrane System: (A) Angled view of system. (B) Bottom up view of system. (C) Side view of system.

## 4 Dependencies

The program is available as a python package [1]. GROMACS [7] must be installed and accessible to your system via the command line `gmx`.

